# SPIN: A Scalable Bioinformatics Pipeline for Screening Pathogenicity-Related Host-Pathogen Protein INteractions Using AlphaFold3

**DOI:** 10.64898/2026.04.21.719732

**Authors:** Zhenghong Bao, Harsh Khanna, Bhuvan Dhand, Wardatou Boukari, Trishna Tiwari, Mousami Poudel, Carlos D. Messina, Mukesh Jain, Jose C. Huguet-Tapia, Rosemary Loria

## Abstract

Traditional experimental approaches have greatly advanced our understanding of protein-protein interactions (PPIs) that govern host susceptibility or resistance. Nonetheless, the molecular characterization of microbial effectors and their cognate host targets remains challenging in many economically important plant pathosystems. AlphaFold3 (AF3) has transformed structural biology by achieving near-experimental accuracy in protein structure and complex prediction. A bioinformatics pipeline SPIN (<u>S</u>creening <u>P</u>athogenicity-Related Host/Pathogen Protein <u>IN</u>teractions) was developed for large-scale prediction of interactions between the host and pathogen-secreted proteins. SPIN integrates three core modules: pathogen and host protein preprocessing using bioinformatics tools (SignalP6.0, OrthoFinder, CD-HIT), AF3-based interaction modeling, supported by automated input generation and output filtering. However, AF3 training bias toward mammalian proteins necessitated careful evaluation in plant systems. Benchmarking against experimentally validated plant PPIs revealed that derivative metrics emphasizing interfacial geometry and residue-level contact (ipSAE and pDockQ) provide superior discrimination compared to native global confidence measures (pTM and ipTM), particularly for proteins with intrinsically disordered regions. A multi-metric confidence scoring framework combining pLDDT, PAE, ipSAE, and pDockQ, improved prediction reliability by enhancing recall and reduced false positives through robust assessment of structural confidence and interface quality. For proof-of-concept, SPIN was applied to examine the molecular landscape underlying two economically important diseases of citrus (*Citrus* L.) caused by *Candidatus* Liberibacter asiaticus and *Ca.* Phytoplasma citri. Both pathogens are phloem-limited and cause distinct symptoms, citrus greening and witches’ broom, respectively. AF3-predicted interactome data revealed conserved host colonization strategies alongside disease-specific molecular mechanisms, demonstrating the utility of SPIN for dissecting and supporting mechanistic studies in plant-pathogen interactions.

## 1 Introduction

Plant immunity against phytopathogens is contingent upon a mélange of host defense responses leading to recalibration of transcriptional and metabolic networks, hormone signaling, and resource allocation. Comprehensive insight into the molecular mechanisms underlying immune cognizance and host susceptibility is crucial for sustainable agriculture and food security (Ryu et al., 2025; Dodds et al., 2010). A critical understanding of how pathogens deploy virulence factors to circumvent plant immunity for host colonization is fundamental to developing durable disease resistance (Ogbuji et al., 2025; Gilbertson et al., 2025; Oliva et al., 2019). Pathogen-associated molecular pattern (PAMP)-Triggered Immunity (PTI) relies on the perception of pathogen-derived elicitors by the cognate cell surface-localized pattern recognition receptors (PRRs) and subsequent defense signaling outputs (Jones and Dangl, 2006). However, plant pathogens often circumvent PTI by delivering effector proteins into the apoplast or directly into host cell cytoplasm to establish a productive colonization and infection cycle (Xiang et al., 2025; Kainat et al., 2025; Zhang et al., 2022; Bialas et al., 2018). Effector-triggered immunity (ETI) is mediated through intracellular coiled-coil nucleotide-binding leucine-rich-repeat (CC-NLR, CNL) and Toll/interleukin-1 receptor nucleotide-binding leucine-rich-repeat (TIR-NLR, TNL) receptors providing robust race-specific resistance against pathogens (Cui et al., 2015). Consequently, the identification of pathogen effectors and cognate host protein interactions is crucial for molecular characterization of virulence mechanisms and host immune signaling pathways (Lovelace et al., 2023) and guiding molecular breeding for disease resistance via transgenic or gene editing workflows (Ogbuji et al., 2025; Oliva et al., 2019).

Mechanistic protein-protein interaction (PPI) studies are crucial for understanding the molecular landscape underlying growth, development, disease resistance/tolerance, and charting large-scale interaction networks in systems biology research. Several biophysical, biochemical, or genetic experimental methods have been developed for experimental validation of host–pathogen protein interactions, including co-immunoprecipitation, yeast two-hybrid, split-luciferase complementation, pull-down (Mocăni ă et al., 2025; Rao et al., 2014), and *in planta* transient expression assays (Berendzen et al., 2012). However, large-scale effector-target screening continues to be costly, labor-intensive, and time-consuming, making it impractical for the entire host/pathogen proteomes. Most importantly, validation of the virulence contribution of each effector protein is contingent upon the functional genomics data which in turn relies on availability of pathosystem-specific molecular tool kits. Genome-wide predictions of the effector arsenal of the pathogen remain challenging due to structural and functional diversity amongst the effectors and microbial secretion mechanisms as reviewed in Lovelace et al. (2023).

*In silico* protein structure prediction methods have been developed as inexpensive and high throughput alternatives to empirical validation with adequate accuracy estimates. Artificial intelligence (AI)–driven platforms such as RoseTTAFold, ColabFold and AlphaFold have transformed protein structural biology and computational biochemistry, enabling unprecedented accuracy in predicting protein structures and interactions (Akdel et al., 2022; Wang et al., 2024; Kim et al., 2025). RoseTTAFold architecture is based on three-track neural network integrating the primary protein sequence, tertiary structures, and interactions in parallel, enabling rapid and accurate protein structure prediction significantly faster than the traditional methods.

By contrast, AlphaFold (Jumper et al., 2021) and its subsequent iterations namely, AlphaFold2 and AlphaFold-Multimer introduced the Evoformer architecture, which analyzes evolutionary multiple sequence alignment (MSA) data to predict complex structural features of single proteins. AlphaFold-Multimer extended this framework to model protein–protein interactions and multimeric assemblies with high accuracy. AlphaFold3 (AF3) builds on these foundations by incorporating the Pairformer module for the pair-weighted MSA averaging, reducing computational memory usage by approximately 30% (Abramson et al., 2024). AF3 also integrates diffusion-based modeling and raw atomic coordinate representations, enabling predictions of the structural dynamics of interacting proteins, homo- or heteromeric complexes, nucleic acids, prosthetic groups, small molecules, and ions within physiological constraints with near-experimental accuracy (Abramson et al., 2024). Despite these advances, computational prediction of host-pathogen protein interactions underlying disease progression remains an arduous task due to the large scale of genomic data, the diversity of scoring metrics that must be interpreted, and the lack of integrated, automated workflows.

To address these challenges, we developed SPIN (<u>S</u>creening <u>P</u>athogenicity-Related Host/Pathogen Protein <u>IN</u>teractions), a scalable bioinformatics pipeline for high-throughput screening of host protein interactions with the secreted pathogen proteins using AF3. SPIN incorporates multiple preprocessing steps-including signal peptide detection for pathogen proteins, orthogroup clustering, and redundancy filtering, followed by AF3-mediated structural interaction modeling. Since the initial AF3 training dataset was largely reliant on mammalian proteins, AF3 interaction scores were validated for experimentally verified ground truth plant pathogen–host protein interactions and a negative-control dataset. A multi-metric scoring framework for confidence ranking incorporated predicted Template Modeling (pTM) and interface predicted Template Modeling (ipTM) scores (Zhang and Skolnick, 2004; Jumper et al. 2021), and structure-confidence and interface-quality metrics, including predicted Local Distance Difference Test (pLDDT), Predicted Aligned Error (PAE), interaction prediction Score from Aligned Errors (ipSAE) (Dunbrack, 2025), and predicted DockQ (pDockQ) (Bryant et al., 2022). While ipSAE prioritizes well-aligned interface residues with down-weighing disordered segments, pDockQ evaluates interface quality based on local confidence and inter-chain contacts. For proof-of-concept, SPIN predictions of host-pathogen protein interactions were examined to understand the molecular mechanisms underlying two important diseases of citrus (*Citrus* L.).

Citrus is one of the highest-value fruit crops in international trade, and Florida has historically been one of the world’s largest citrus producers (Graham et al., 2024). Citrus greening (Huanglongbing or HLB) is arguably the single most destructive citrus disease worldwide. HLB is associated with the gram-negative, hitherto uncultured, and phloem-limited α-Proteobacterium *Candidatus* Liberibacter asiaticus (CLas) in Asia and the Americas (Bove, 2006) and transmitted by the Asian citrus psyllid (ACP) *Diaphorina citri* Kuwayama (Gottwald et al., 2010). *Ca.* L. americanus (CLam) and *Ca.* L. africanus (CLaf) are also associated with citrus greening disease in Brazil and Africa, respectively. Reactive Oxygen Species (ROS) mediated oxidative stress, phytohormone imbalance, and callose and starch deposition result in impaired nutrient transport and assimilate partitioning, phloem necrosis, canopy and root loss, and eventual decline of the infected tree (Ma et al. 2022). Incidence of slow (progressive) or sudden citrus decline disease (CDD) is increasing and often attributed to mixed infections of CLas and *Ca.* Phytoplasma citri strains 16SrII-B and -C (Noorizadeh et al. 2022). Akin to CLas, *Ca.* P. citri 16SrII-B and -C strains are intracellular and phloem-limited phytopathogens transmitted by the leafhopper *Neoaliturus haematoceps* and ACP (Zreik et al. 1995). *Ca.* Phytoplasma infections result in witches’ broom symptoms such as aberrant branching architecture and lack of reproductive transition thereby compromising fruit yield.

Liberibacters (Duan et al. 2009, Prasad et al. 2016) and Phytoplasmas (Oshima et al. 2013) lack the canonical effector secretion systems and rely exclusively on Sec-dependent translocation for secreting effector proteins directly into the host cell cytoplasm, evading host defenses, and manipulate host metabolism for an intracellular lifestyle. Pathogenicity-related contributions of several of these small-secreted proteins and/or hypothetical proteins remain sparsely understood. We applied the bioinformatics pipeline SPIN to gain structural insights into these host-pathogen protein interactions in *Ca.* Liberibacter- and *Ca.* Phytoplasma-infected citrus phloem (Fig. 1). The primary objective was to assess whether AF3-based predictions could (a) provide insights into molecular mechanisms of disease progression, (b) support downstream experimental workflows by enabling informed prioritization of biologically relevant candidate interactions, and (c) if the computational workflow is robust enough to compare and contrast the infection strategies of two discrete pathogens occupying the same niche for colonization.

**Fig. 1:**
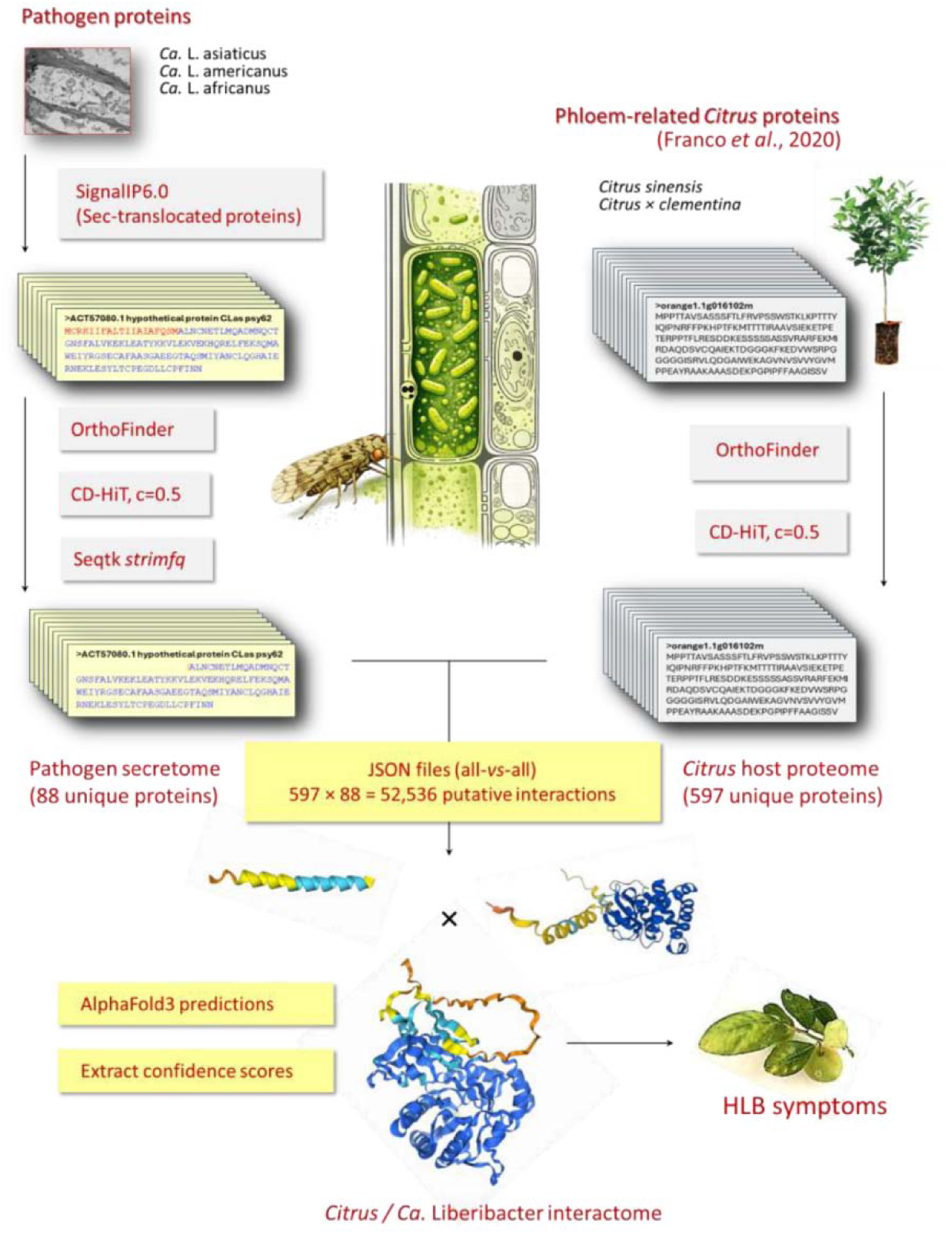
Schematic of SPIN pipeline for high throughput prediction of host-pathogen protein interactions. The SPIN pipeline grouped the *Ca.* Liberibacter secretome into 88 unique proteins which were subsequently paired with 597 unique *Citrus* phloem-associated proteins. Pathogenicity-related 52,536 likely interactions were structurally analyzed via AlphaFold3. Likewise, 14 unique *Ca.* Phytoplasma citri proteins were also interrogated with *Citrus* proteins for 8,358 likely interactions.

## 2 Materials and methods

### 2.1 AlphaFold3 benchmark datasets for parameter selection and performance validation

We used a set of positive controls that consist of experimental ground truth interactions extracted from the Protein Data Bank (https://www.rcsb.org/) and the pathogen-host interaction PHI-base (http://www.phi-base.org/). For comparison and AF3 validation, a negative control dataset of protein pairs was used comprising of protein pairs with no evidence of biological interactions based on Gene Ontology (GO) (https://geneontology.org/) and Kyoto Encyclopedia of Genes and Genomes (KEGG) (https://www.genome.jp/kegg/pathway.html) pathway annotations. Proteins were paired only if they belonged to distinct and functionally unrelated biological processes or pathways, with no overlap in GO biological process terms or KEGG pathway membership. Additionally, any proteins previously associated with host-pathogen interactions were excluded to ensure that the negative set reflected biologically implausible interactions. In total 80 ground-truth positive protein interaction pairs and 324 negative (non-interacting) protein pairs were curated (Suppl. Table S0).

### 2.2 Host (*Citrus* phloem-related) and pathogen (*Ca.* Liberibacter and *Ca.* Phytoplasma citri) protein datasets

Protein sequences encoded by 18 fully assembled genomes of three HLB-associated Liberibacter species (CLas, CLam and CLaf) and eight *Ca.* P. citri strains were retrieved from NCBI RefSeq and GenBank databases. Details for each genome, including accession numbers and strain identifiers, are provided in Suppl. Table S1. These datasets included 18,396 *Ca.* Liberibacter and 2,812 *Ca.* P. citri proteins that were the starting point of the analysis. The enriched phloem-related proteome of Washington Navel sweet orange [*C. sinensis* (L.) Osbeck] (Franco et al. 2020) provided the host repertoire for downstream interaction prediction with *Ca.* Liberibacter and *Ca.* Phytoplasma secreted proteins. Full-length sequences of 733 phloem-related *Citrus* proteins were retrieved from the annotated *C. sinensis* v1.1 and *C. clementina* v1.0 genomes (https://phytozome-next.jgi.doe.gov/) (Suppl. Table S2).

### 2.3 Computational workflow for host-pathogen protein processing and AF3-mediated prediction of protein-protein interactions

This computational workflow consisted of three major steps that generate pathogen and host protein datasets for AF3-mediated interaction modeling. The procedure integrates secretion prediction, orthogroup inference, clustering-based refinement, and standardized host-pathogen protein pairing to ensure consistent preprocessing and reproducible generation of input files for downstream AF3 multimer predictions.

#### 2.3.1 Pathogen and host protein processing

Proteins within the same orthogroup often share functions and are likely to interact or participate in similar pathways, providing insights into infection mechanisms and immune evasion. A hybrid strategy combining phylogenomic and sequence-based approaches was employed to classify, refine, and analyze protein sets from the plant host and associated pathogen. SignalP 6.0 (Teufel et al., 2022) was used to identify signal peptides from pathogen proteins to determine their secretion potential. OrthoFinder v2.5.5 (Emms and Kelly, 2019) was used to group secreted proteins into orthogroups based on gene-tree reconciliation and species-tree topology to preserve the clusters representing speciation-derived evolutionary relationships. A second orthogroup clustering step with CD-HIT v4.6.8 (50% identity threshold) (Fu et al., 2012) was deployed to reduce redundancy. This step separated highly divergent proteins within the same orthogroup, potentially revealing sub-functionalization or lineage-specific adaptations, while simplifying downstream analyses and reducing computational load. CD-HIT provides flexibility in clustering stringency by allowing user-specified identity thresholds (e.g., 90% or 50%), Signal peptides were removed from the selected proteins using the *strimfq* command from the Septik toolkit v1.4. Similarly, host phloem-related proteins were first processed using OrthoFinder and subsequently clustered at a 50% sequence identity threshold using CD-HIT.

#### 2.3.2 Host-pathogen protein pairing and structure predictions

The host and pathogen proteins sequences were converted into AF3-compatible JSON (JavaScript Object Notation) input files using an *all-versus-all* pairing strategy. Each JSON file contained up to a user-specified number of pathogen-host protein pairs (default 30) with optional parameters allowing specification of copy numbers for each protein chain. Structural predictions were generated using a local installation of AF3 v3.0.0) deployed as a Singularity image on a High-Performance computing cluster (HiPerGator; University of Florida) leveraging an Ampere A100 GPU (80 GB), 8 CPU cores, and 120 GB of memory, with CUDA Toolkit 12.4 environment and GPU driver version 550.127.05. The installation utilized the officially released GitHub version, publicly requestable model weights, and the corresponding database snapshot (downloaded in January 2025). All runs were executed under standardized settings to ensure reproducibility across large-scale screening experiments. Predictions generated from the local installation were processed through an automated post-analysis pipeline to extract confidence metrics, including ipTM, pTM, ipSAE, and pDockQ scores. ipTM and pTM are native AF3 confidence scores, whereas ipSAE and pDockQ were computed post hoc with the open-source tool at github.com/dunbracklab/IPSAE. These metrics were compiled into structured CSV output files for downstream ranking and comparative analyses.

For comparison and cross-platform validation, selected protein interaction pairs were also submitted remotely to the AlphaFold server (alphafoldserver.com/) operating within a managed production environment. Structure predictions obtained from AlphaFold server were processed individually using a custom Python script (post_analysis_AF3server.py; https://github.com/baozhuf/SPIN) to extract the same confidence metrics for comparison.

#### 2.3.3 Integrated structural and interface confidence metric scores for AF3-predicted protein complex evaluation

Predicted protein complexes generated by AF3 were evaluated using the native model confidence outputs for the global structural confidence (pTM and ipTM) and residue-level reliability (pLDDT and PAE matrix). Additionally, two additional interface-focused metrics (ipSAE and pDockQ) were computed from the predicted structures (Table 1). The ipSAE score was calculated by averaging confident inter-chain residue pairs derived from the PAE matrix, providing an estimate of interaction confidence based on inter-chain alignment errors. In parallel, pDockQ was computed by integrating interface pLDDT values with inter-chain contact density to estimate the likelihood of a correct docking interface. Global metrics ensured overall structural stability of the predicted complex, while interface metrics evaluated the reliability and physical plausibility of the predicted interaction surface. The combined criteria were designed to reduce false-positive predictions arising from models with high global confidence but weak or poorly supported interfaces. These thresholds were applied uniformly across all large-scale host-pathogen interaction screening analyses.

**Table 1:** Native and derived (post hoc computed) confidence scores in AlphaFold3 predictions of protein interaction complexes.

| Confidence score | What it measures | Score range | Interpretation |
| --- | --- | --- | --- |
| pLDDT<br>(predicted Local Distance Difference Test) | Estimates local structural accuracy; provides per-residue confidence scores | 0-100 | > 90: Highly reliable<br>70–90: Generally reliable backbone prediction |
| PAE<br>(Predicted Aligned Error) | Measures confidence in relative position and orientation between residue pairs | 0-30 Å | < 5 Å: High confidence<br>5–15 Å: Moderate confidence |
| pTM<br>(predicted Template Modeling) | Assesses accuracy of overall 3D structure and global fold | 0-1 | > 0.8: High confidence<br>> 0.5: Correct overall backbone prediction |
| ipTM<br>(interface predicted Template Modeling) | Evaluates relative positioning and orientation between chains in a complex | 0-1 | > 0.8: High confidence<br>0.6–0.8: Intermediate/grey area |
| ipSAE<br>(interface score from Aligned Errors) | Considers only thr residue pairs with PAE below a cutoff (typically 10 Å), reduces effects of disordered regions | 0-1 | > 0.6: General cutoff for confident interactions |
| pDockQ<br>(predicted DockQ) | Estimates interface quality using inter-chain contacts and average interface pLDDT | 0-1 | > 0.5: High quality<br>> 0.23: Acceptable quality |

Protein-protein interaction classification was performed using predefined thresholds for each metric. Positive interactions were defined using a combined evaluation of global structural confidence (ipTM ≥ 0.7 and pTM ≥ 0.5) and interface-focused metrics (ipSAE ≥ 0.6 and pDockQ ≥ 0.23). Threshold values were optimized using the benchmark dataset described above, using standard statistical measures including precision, recall, F1-score, and false positive rate (FPR). A protein pair was classified as an interaction when both global structural confidence and interface quality exceeded the optimized cutoff values.

## 3 Results

### 3.1 AlphaFold3 accurately predicted and separated verified plant protein interactions from poor interactions

Although the training dataset spans diverse taxa (Abramson et al. 2024), experimentally resolved plant protein structures remain comparatively underrepresented, which may affect predictive performance in plant systems, particularly for less conserved protein families. SPIN is built around AF3 as its central prediction module. To evaluate AF3 prediction performance, we curated a benchmark dataset comprising 80 experimentally validated plant protein-protein interactions (ground-truth PPIs) and 324 plant protein pairs comprising of biologically implausible ground-truth non-interacting proteins that are excluded by any GO and KEGG pathway associations across molecular function, cellular component, or biological process categories (Suppl. Table S0). This curated dataset was analyzed twice using both the locally (HiPerGator) (Fig. 2A, B) and remotely (alphafoldserver.com/) deployed AF3 (Fig. 2C, D) and the resulting ipTM, pTM, ipSAE, and pDockQ scores were extracted (Suppl. Table S0).

**Fig. 2:**
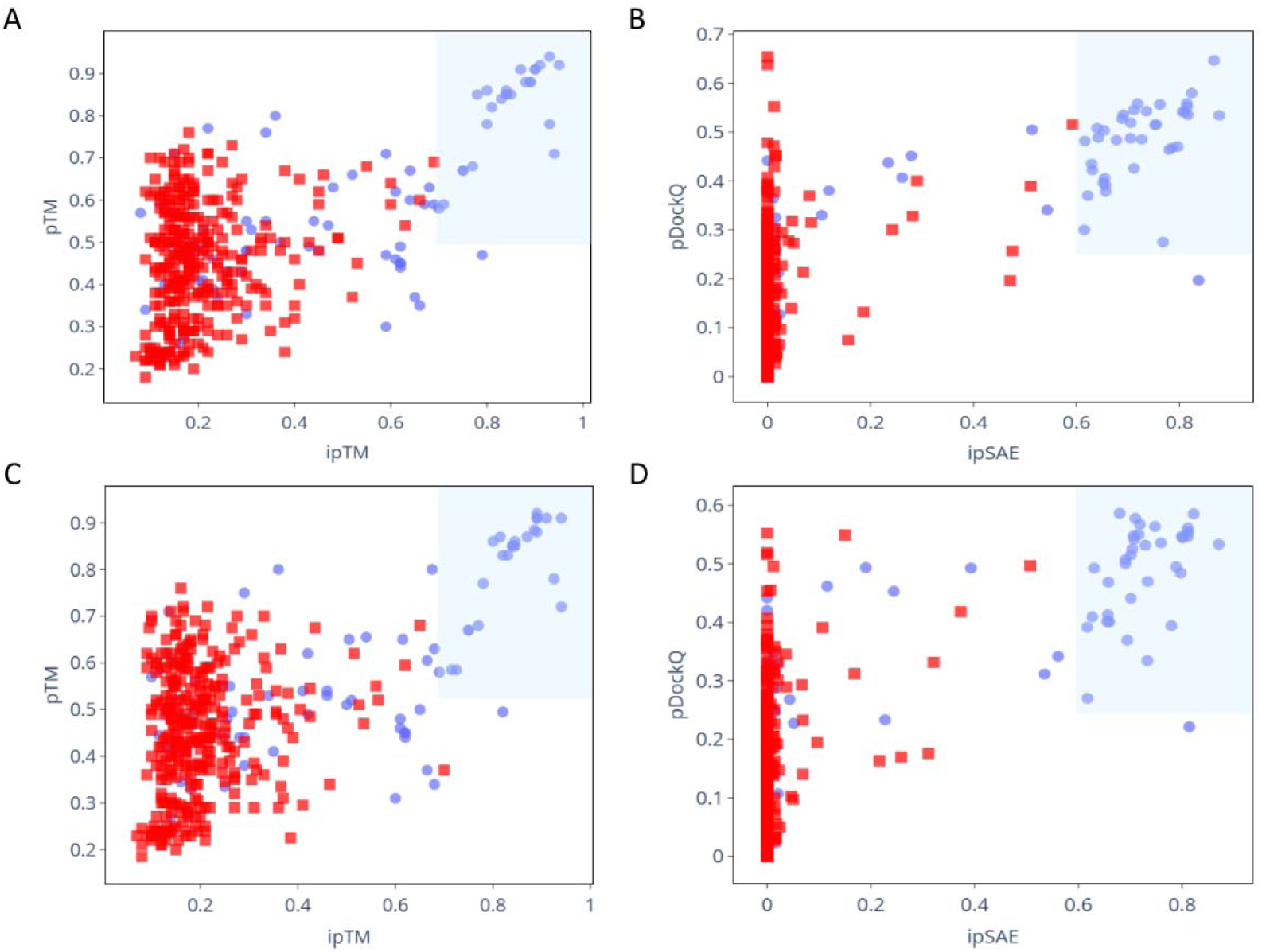
AlphaFold3 predictions for the empirically validated plant protein-protein interactions. **A, C:** Scatter plot of ipTM and pTM scores (thresholds 0.7 and 0.25, respectively) for 80 experimentally verified protein interactions 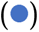 and 324 protein pairs expected to interact poorly 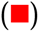. **B, D:** Scatter plot showing efficient separation of ground-truth positive 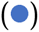 and ground-truth negative 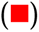 interactions based on the ipSAE-pDockQ scores (thresholds 0.6 and 0.23, respectively). These curated datasets were analyzed twice using both the locally (HiPerGator) **(A, B)** and remotely (alphafoldserver.com/) deployed AlphaFold3 **(C, D).** By contrast to ipTM-pTM metric pair, the ipSAE-pDockq confidence scores improved the recall of true positives from 25 to 37 **(A, B)** and from 24 to 35 **(C, D)** and the F1 score from 0.48 to 0.63 **(A, B)** and 0.46 to 0.61 **(C, D)** for previously validated protein interactions.

Figure 2A represents the distribution of ipTM and pTM scores for the 80 ground-truth PPIs and 324 non-interacting protein pairs (HiPerGator run). All false positives were eliminated (FPR = 0) at a combined threshold of ipTM = 0.7 and pTM = 0.5, yielding 100% precision but only 30% recall (Table 2). This stringent cutoff ensured high confidence in predicted interactions but 55 of 80 true PPIs were not included within the decision boundary, thus reducing the sensitivity. The corresponding F1 score = 0.48 reflected this trade-off. To improve the recall without compromising precision, interface-specific derived metrics ipSAE and pDockQ were also evaluated for the same dataset. Compared to the global ipTM and pTM scores, the combined ipSAE and pDockQ scores provided improved separation of ground-truth PPIs, and most ground-truth non-PPIs clustered near ipSAE ≈ 0 (Fig. 2B). At the ipSAE = 0.6 and pDockQ = 0.23 thresholds, an increased 46% recall was achieved while maintaining 100% precision and FPR = 0 (Table 2). The F1 score improved to 0.63, highlighting the advantage of incorporating interface-focused metrics for adequate and accurate detection of true PPIs.

**Table 2:** Summary of the classification performance of AlphaFold3 using global (ipTM, pTM) and interface-focused (ipSAE, pDockQ) confidence thresholds across local (HiPerGator) and remote (alphafoldserver.com/) environments.

| Computing environment | AF3 confidence score thresholds | Classification statistical metrics* |  |  |  |  |  |
| --- | --- | --- | --- | --- | --- | --- | --- |
|  |  | TP | FN | Precision | Recall | F1 | FPR |
| HiPerGator | ipTM =0.7, pTM =0.5 | 25 | 55 | 1 | 0.31 | 0.48 | 0 |
|  | ipSAE =0.6, pDockQ =0.23 | 37 | 43 | 1 | 0.46 | 0.63 | 0 |
| alphafoldserver.com/ | ipTM =0.7, pTM =0.5 | 24 | 56 | 1 | 0.30 | 0.46 | 0 |
|  | ipSAE =0.6, pDockQ =0.23 | 35 | 45 | 1 | 0.44 | 0.61 | 0 |

\*TP: true positives; FN: false negatives; FPR: false positive rate; Precision = $TP/(TP+FP)$ ; Recall (sensitivity) = $TP/(TP+FN)$ ; F1: harmonic mean of precision and recall

The benchmark datasets of ground-truth PPIs and non-interacting protein pairs were also evaluated remotely at alphafoldserver.com/ (Figs. 2C, D). However, slight variations were discernible in the ipTM, pTM, ipSAE, and pDockQ scores for the evaluated protein pairs across the two platforms (also see Table 3). Nevertheless, interface prediction scores (ipSAE = 0.6 and pDockQ = 0.23) improved the recall and F1 score (0.44 and 0.61, respectively) while maintaining 100% precision and FPR = 0 (Table 2). Multi-metric scoring integrating ipTM, pTM, ipSAE, and pDockQ is essential for evaluating predicted host-pathogen protein interactions in high-throughput screens, particularly in non-model systems. In addition, selecting appropriate confidence thresholds that yield zero false positives helps avoid wasteful downstream experimental validation.

**Table 3:** Reproducibility of AlphaFold3 confidence metrics across repeated runs and computational platforms. Absolute differences (Δ) were calculated between repeated runs using identical random seeds (2025). Values represent the number of protein pairs (n = 404) exhibiting a metric difference ≥0.2.

| Comparison | $\Delta_{ipTM} \geq 0.2$ | $\Delta_{pTM} \geq 0.2$ | $\Delta_{ipSAE} \geq 0.2$ | $\Delta_{pDockQ} \geq 0.2$ | Avg* | %* |
| --- | --- | --- | --- | --- | --- | --- |
| HPG (run1 vs. run2) | 0 | 0 | 0 | 0 | 0 | 0 |
| Server (run1 vs. run2) | 2 | 0 | 2 | 7 | 2.8 | 0.6 |
| Mean (Server-HPG) | 12 | 1 | 8 | 19 | 10 | 2.2 |

\*Avg represents the average number of pairs exceeding the threshold across all four confidence metrics evaluated; % indicates the ratio of the average count relative to the total number of protein pairs evaluated.

AF3 prediction outputs for two contrasting protein pairs are shown in Fig. 3, illustrating these multi-metric scores from a structural perspective. Yeast-two-hybrid screen had previously confirmed biological interaction between the rice blast fungus *Pyricularia oryzae* (syn. *Magnaporthe oryzae*) strain B157 effector (Chain I, APikL2F) (7NMM_I) and *Setaria italica* Heavy Metal Associated domain (HMA)-containing protein (sHMA94) (7NMM_A) (Bentham et al., 2021). AF3 corroborated a high-confidence APikL2F-sHMA94 interaction complex *in silico*, with low PAE both within and between the interacting subunits yielding high ipTM (=0.91), pTM (=0.91), ipSAE (=0.82) and pDockQ (=0.58) scores (Fig. 3A). By comparison, the sugar beet cyst nematode *Heterodera schachtii* effector (ABY49997.1) is not expected to interact with *Arabidopsis thaliana* pectin methylesterase 3 (AT3G14310). AF3 output predicted confident individual protein structures (pTM = 0.62), but high PAE values indicating inter-chain uncertainty in the relative positioning of the two proteins. The overall ipSAE and pDockQ scores (=0.0 and 0.08, respectively) further contradicted any biological interaction under physiological conditions (Fig. 3B).

**Fig. 3:**
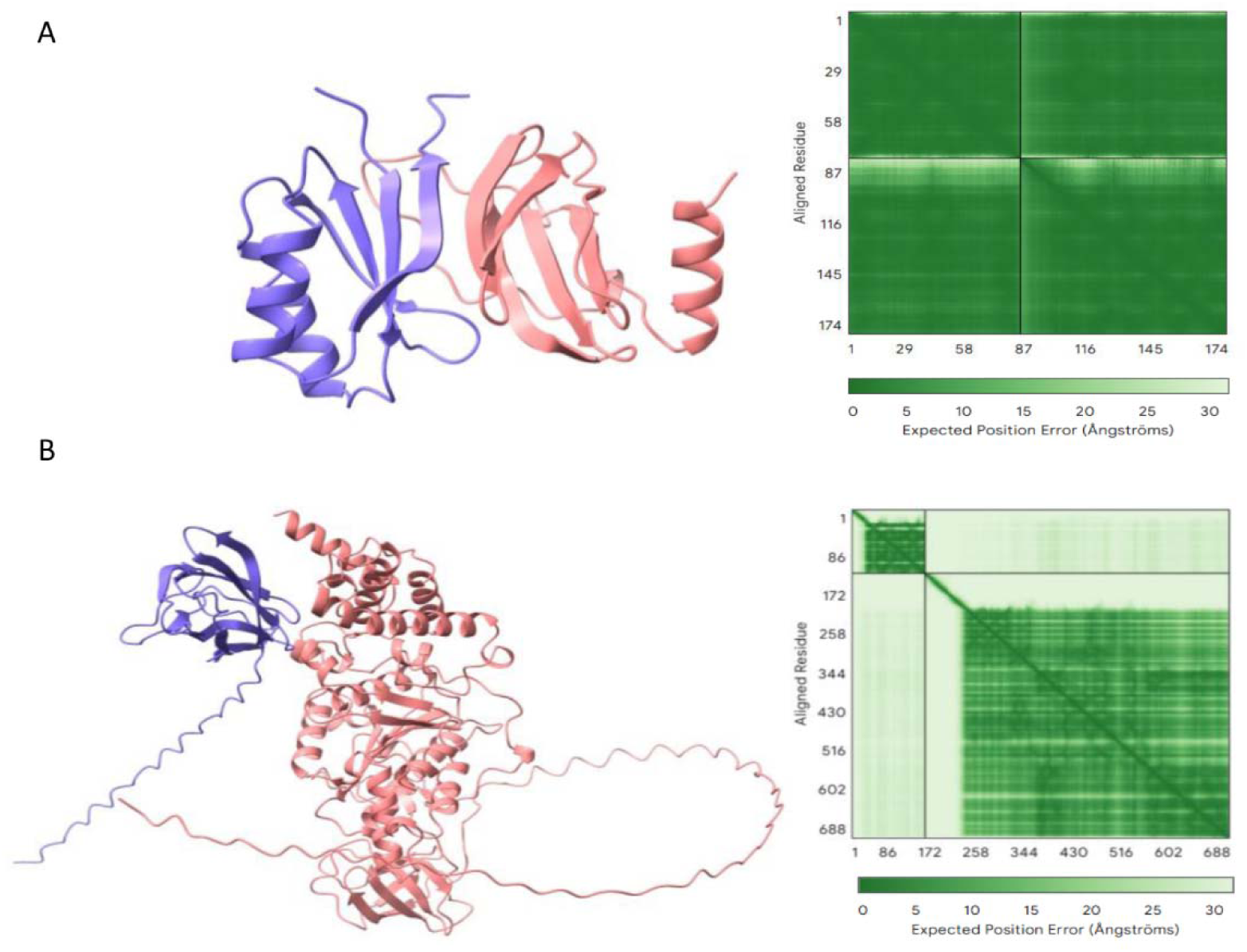
AlphaFold3 interface outputs illustrating predicted interaction models, and PAE matrices showing relative positions of residue pairs within the two interacting protein partners. **A:** AlphaFold3-predicted (and experimentally verified) interaction complex of rice blast fungus *Magnaporthe oryzae* effector (APikL2F) with the *Setaria italica* Heavy Metal-Associated domain (HMA)-containing protein (sHMA94) yielded high ipTM (=0.91), pTM (=0.91), ipSAE (=0.82) and pDockQ (=0.58) scores. **B:** Unlikely interaction between the sugar beet cyst nematode *Heterodera schachtii* effector (ABY49997.1) and *Arabidopsis thaliana* pectin methylesterase 3 (AT3G14310). High AlphaFold3 PAE values indicating inter-chain structural uncertainty with poor ipSAE and pDockQ scores (=0.0 and 0.08, respectively).

#### 3.1.1 Reproducibility and veracity of AlphaFold3 predictions across computational environments

To assess repeatability of large-scale AF3 predictions, we evaluated AF3 output data across computational environments using 404 benchmark protein pairs (including both experimentally verified and poor/unlikely interactions). Two independent runs were performed both on the local HiPerGator and on the remote AlphaFold server, all using an identical random seed. Absolute differences greater than 0.2 were quantified for all the four scoring metrics, viz. ipTM, pTM, ipSAE, and pDockQ (Table 3). Within the HiPerGator environment, no metric exceeded the 0.2 threshold between independent runs, indicating complete reproducibility under fixed seed conditions. By contrast, the AlphaFold server yielded minor run-to-run variability. Absolute differences ≥0.2 were observed in two instances for ipTM, 2 for ipSAE, and seven for pDockQ, corresponding to 0.6% of the evaluated interactions.

Cross-platform comparisons for the computing runs across the two environments showed a slightly higher level of discrepancy. Differences ≥0.2 were observed in 12 protein pairs for ipTM, 1 for pTM, 8 for ipSAE, and 19 for pDockQ, corresponding to approximately 2.2% of the dataset (Table 3). These results demonstrate overall reliability of AF3 predictions, with limited variability detected primarily in cross-platform comparisons. The observed differences likely stem from variations in MSA generation (https://github.com/google-deepmind/alphafold3AF3/issues/492), GPU architecture, and floating-point precision rather than stochastic processes, as all runs used identical random seeds.

### 3.2 SPIN: High throughput prediction of host-pathogen protein interactions using AlphaFold3

Interactions between the secreted pathogen proteins (virulence factors/effectors) and the host proteins are essential for facilitating pathogen colonization and immune evasion, thus leading to symptom development and disease progression. Applicability of SPIN integrating bioinformatic tools and AF3 for high throughput prediction of host-pathogen PPIs was examined to understand the host-pathogen ‘*interactome*’ and molecular roadmap underlying greening and witches’ broom diseases of citrus. Both citrus pathogens, *Ca.* Liberibacter and *Ca.* P. citri, are phloem-limited, unculturable and lack the canonical effector secretion systems therefore intractable to traditional functional genomic studies.

#### 3.2.1 Prediction of *Ca.* Liberibacter-secreted protein targets in citrus phloem

Eighteen fully assembled genomes from the three HLB-associated Liberibacters (CLas, CLam, and CLaf) encode a total of 18,396 proteins that were used as input for the SPIN pipeline, together with the citrus dataset consisting of 733 phloem-related proteins (Franco et al., 2020).

From the Liberibacter genomes, SignalP analysis predicted 798 Sec-translocated proteins (Suppl. Table S3). Notably, several Sec-translocated Liberibacter proteins have relatively small size (average 263 amino acid residues) and some of them have been annotated as hypothetical proteins (HPs) lacking any conserved structural or functional Pfam domains. Despite a highly reduced genome, a total of 142 proteins were identified as HPs within the Sec-translocated proteome of *Ca.* Liberibacters underscoring their likely role in HLB progression. The SPIN pipeline grouped the Liberibacter secretome into 88 unique proteins (45 HPs) (Suppl. Table S4), which were then paired with the unique 597 *Citrus* phloem-associated protein (Suppl. Table S5) dataset generating 52,536 pairs for the putative protein complexes. The resulting pairs were organized into 597 JSON files (*Citrus vs.* Liberibacter proteins) and executed on the HiPerGator cluster and the AlphaFold server. The screened protein interaction pairs had an average combined sequence length of 679 amino acid residues (Suppl. Table S6). The average prediction time was ∼14 minutes per interaction, with the average CPU utilization of 49% (range: 44%–54%) and an average memory efficiency of 35% (ranging from 27% to 87%).

The use of JSON files optimized pipeline execution and enabled efficient, high-throughput prediction of host-pathogen protein interactions. The density distribution of AF3 native confidence scores (ipTM and pTM) for the 52,536 predicted PPIs (Suppl. Table S7) is presented in Fig. 4A. Five protein pairs are missing from Table S7 because their amino acid sequences are exceptionally long, for example WP_012778561.1-VS-Ciclev10007222m (3994 aa), which prevented us from running AF3 predictions on either HiPerGator or the AlphaFold server. AF3-mediated structural modeling predicted highly confident 82 PPIs (pTM ≥ 0.5, ipTM =0.8-1.0), whereas moderate (pTM ≥ 0.5, ipTM =0.6-0.8) to low (pTM ≥ 0.5, ipTM =0.3-0.6) confidence scores were observed for 924, and 6,167 paired interactions, respectively. SPIN analysis (Fig. 4A) corroborated the previously identified Sec-translocated CLas effectors and their cognate citrus protein targets consistent with the previously published yeast-two-hybrid and co-immunoprecipitation data (Table 4).

**Fig. 4:**
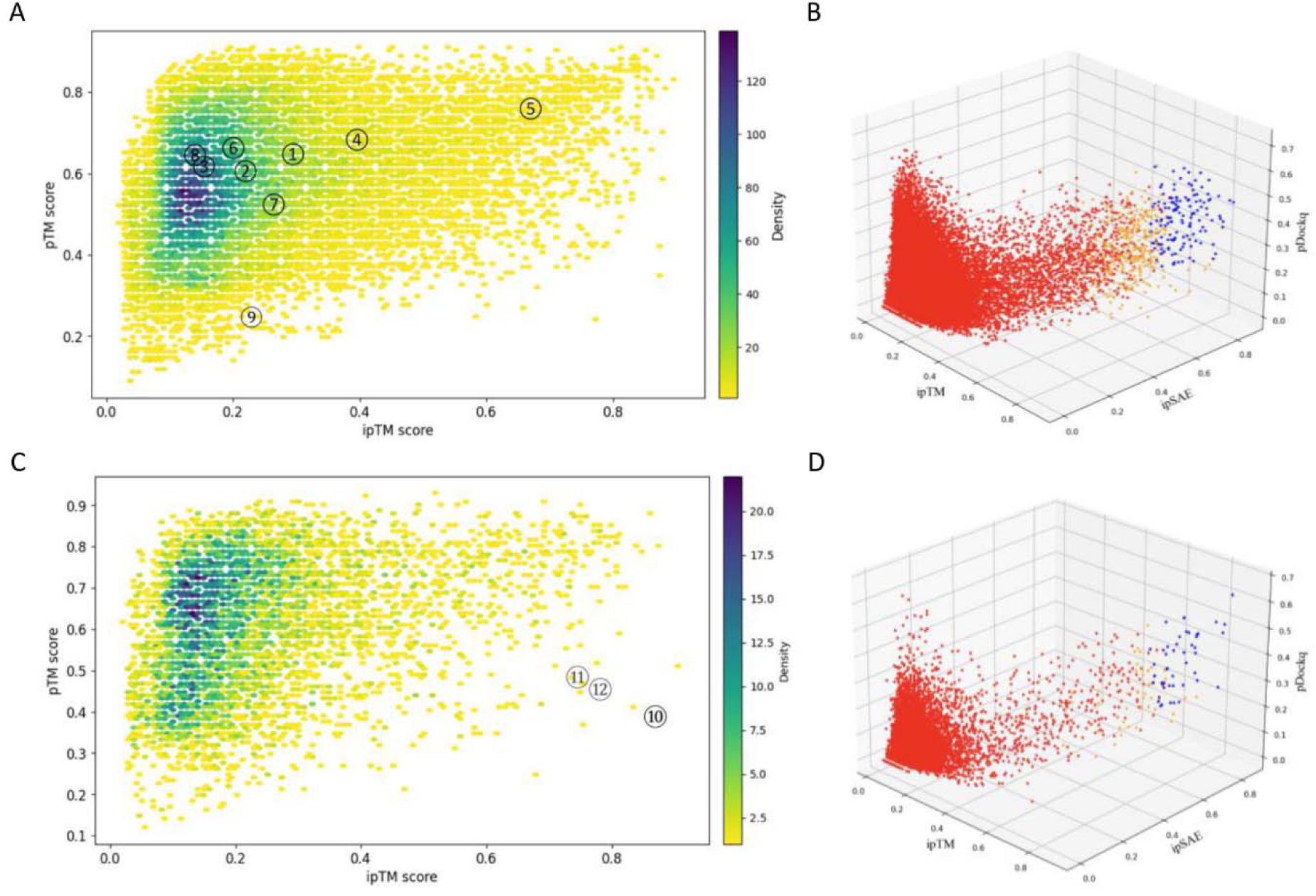
High throughput AlphaFold3 predictions of pathogenicity-related protein interactions between *Citrus* phloem-related proteins and secreted proteins of **(A, B)** *Ca.* Liberibacter and **(C, D)** *Ca.* Phytoplasma citri. **A, C:** Density distribution of AlphaFold3 confidence metrics (ipTM and pTM) for 52,536 predicted *Citrus*-*Ca.* Liberibacter and 8,358 *Citrus*-*Ca.* P. citri protein interactions. **B, D:** 3D scatter plot classifying protein interactions based on pTM, ipSAE and pDockQ scores: interactions with low ipTM score (<0.6) are shown in red 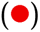, with adequate ipTM scores (>0.6) but scoring low on ipSAE (<0.6) or pDockQ (<0.23) scores are shown in orange 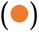. The high-confidence protein interactions (ipTM >0.6, ipSAE >0.6, pDockQ >0.23) are marked in blue 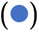 and may be pursued for further biological validation (136 for *Ca.* Liberibacter, and 50 for *Ca.* P. citri, respectively). SPIN data corroborated previously validated interactions of *Ca.* Liberibacter **(A)** and *Ca.* P. citri **(C)** with *Citrus* proteins (marked 1-9, and 10-12, respectively) (also see Table 4).

**Table 4.**
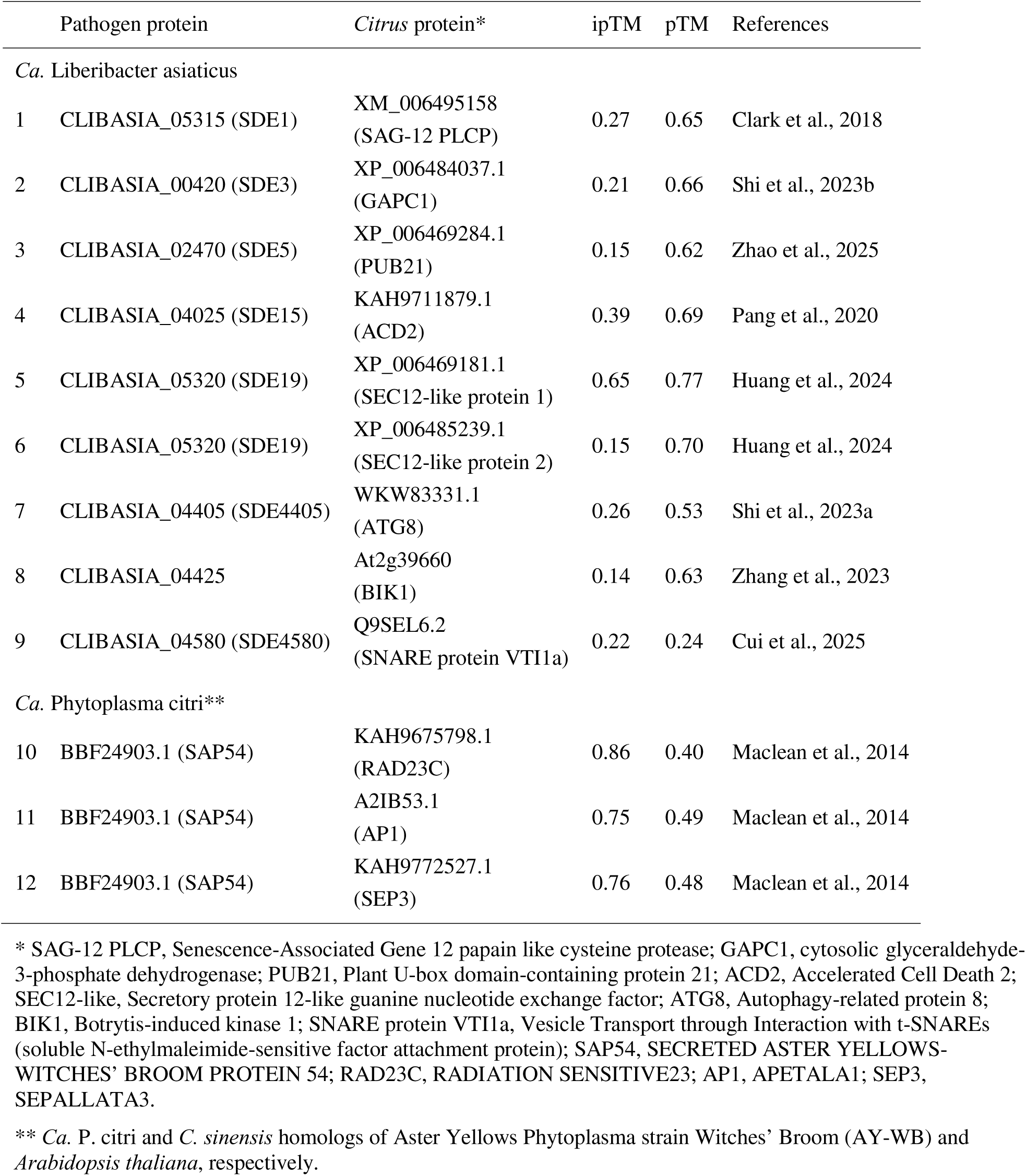
Native AlphaFold3 confidence scores for the experimentally validated interaction complexes of *Citrus sinensis* phloem-related proteins with the secreted proteins of *Ca*. Phytoplasma citri.

As previously demonstrated, ipSAE and pDockQ scores show superior discriminatory power in segregating experimentally verified PPIs and non-interacting partners (Fig. 2). Therefore, *Citrus*-Liberibacter PPIs were further filtered through a three-stage confidence pipeline (ipTM ≥ 0.6, ipSAE ≥ 0.6 and pDockQ ≥ 0.23) to separate interactions with structurally reliable interface geometry yielding 136 and 292 PPIs with high to moderate confidence, respectively (Fig. 4B). Without the second-stage filtering using ipSAE and pDockQ, interactions with adequate ipTM scores, but poor local confidence and interchain contacts would have been incorrectly advanced to downstream analysis, resulting in wasteful experimental overhead. A stringent, two-stage filtering is therefore pragmatic to identify targets for effector proteins in a high throughput screen.

#### 3.2.2 Prediction of *Ca.* Phytoplasma citri-secreted protein targets in *Citrus* phloem

Six fully assembled and annotated *Ca.* P. citri genomes (Suppl. Table S1) collectively encode 2,812 proteins, with 53 proteins containing signal peptides for Sec-dependent translocation (Suppl. Table S8). The Sec-translocated proteins were grouped into 14 distinct orthogroups, yielding 14 unique proteins with an identity threshold of 0.5 in CD-HIT (Suppl. Table S9). Native AF3 structural prediction confidence scores (pTM and ipTM) for 8,358 candidate PPIs (14 *Ca.* P. citri × 597 *Citrus* phloem-related proteins) are shown in Fig. 4C. Based on the prerequisite for a confident overall structure prediction (pTM ≥ 0.5), AF3-mediated structural modeling predicted 16 PPIs with high (ipTM =0.8-1.0), 201 with moderate (ipTM =0.6-0.8), and 1,042 with low (ipTM =0.3-0.6) confidence scores (Suppl. Table S10). Furthermore, three-stage confidence filtering (ipTM ≥ 0.6, ipSAE ≥ 0.6 and pDockQ ≥ 0.23) segregated 50 and 49 *Citrus*-Phytoplasma PPIs with high to moderate confidence in interface geometry, respectively (Fig. 4D).

Molecular mechanisms underlying phloem colonization by Phytoplasma and regulation of plant morphogenesis are sparsely understood. Aster Yellows Phytoplasma strain Witches’ Broom (AY-WB) effector SECRETED AY-WB PROTEIN 54 (SAP54) interacts with the RADIATION SENSITIVE23 family proteins (RAD23). RAD23-marked proteasome-mediated degradation of floral development regulators SEPALLATA3 (SEP3) and APETALA1 (AP1) causes homeotic transformation in *A. thaliana* flowers (Maclean et al., 2014). Corroborating these data, SPIN pipeline identified high-confidence interactions between the *Ca.* P. citri SAP54 homolog (BBF24903.1) with the citrus RAD23C, AP1, and SEP3 homologs (KAH9675798.1, A2IB53.1, and KAH9772527.1, respectively) (Fig. 4C, Table 4).

## 4. Discussion

Computational protein modeling and pretrained language models have accelerated the study of effector biology and host-pathogen interactions. However, wider application of structural biochemistry in plant pathology has only recently begun to emerge. AlphaFold-Multimer screen of tomato (*Solanum lycopersicum*) identified apoplastic P69B subtilase as the target hub for 1879 effectors secreted by seven diverse bacterial, fungal and oomycete pathogens (Homma et al., 2023). AlphaFold-Multimer screening of the soybean root apoplast proteome identified cysteine- and serine-protease inhibitor complexes associated with charcoal rot disease caused by *Macrophomina phaseolina*, revealing novel targets for improved disease resistance (Prakash et al., 2025).

### 4.1 Enhanced prediction of protein interactions using multi-metric AlphaFold3 scoring

AF3 leverages large-scale sequence and structural databases spanning 350,000 diverse sequences across app. 47 key taxa from Uniclust30 (Mirdita et al., 2017) to infer evolutionary and co-evolutionary constraints (https://deepmind.google/technologies/alphafold) (Abramson et al., 2024). Experimentally resolved plant protein structures remain markedly underrepresented in the AF3 training dataset, comprising only 2,569 proteins, predominantly from the model species *Arabidopsis thaliana*. Notably, less than 1% of protein sequences from major crop species have been structurally characterized (Ceasar and Ebeed, 2024). Calibration of AF3-derived structural and interfacial priors may be suboptimal for plant systems, underscoring the need for cautious interpretation and integrative validation when applying these predictions to plant-pathogen interaction studies.

When evaluated against the ground-truth set of plant PPIs, AF3 exhibited a systematic bias characterized primarily by reduced sensitivity rather than indiscriminate prediction error (Fig. 2). Conservative structural modeling of plant-pathogen PPIs favors higher confidence thresholds at the expense of missing a subset of true interactions. Derivative metrics emphasizing interface geometry and residue-level contact confidence, ipSAE and pDockQ, provide superior discrimination compared to native global confidence measures such as pTM and ipTM (Fig. 2) (Bryant et al., 2022; Zhu et al., 2023).

The reproducibility of AF3 predictions are highly stable within a controlled computational environment (Table 3). Absence of run-to-run variation observed on the local HiPerGator installation indicating deterministic behavior under fixed seed conditions, an essential attribute for large-scale benchmarking studies. By contrast, minor variability on the AF3 Server and a 2.2% cross-platform discrepancy suggested that factors beyond seed control, such as GPU architecture, numerical precision, or backend implementation, can influence confidence metrics in a small subset of cases. As identical seeds were used, these differences likely reflect variation in MSA generation, database versions, or hardware-dependent computations. Although AF3 reduces reliance on deep MSAs, evolutionary information remains central to its modeling framework (Jumper et al., 2021; Peng et al., 2025). Small differences in MSA composition may influence interface confidence, particularly in borderline cases marked by limited phylogenetic data and/or homolog diversity. While high-confidence predictions for protein complexes remain stable, marginal cases are more sensitive to input variation, highlighting a potential reproducibility concern across platforms and pipelines, especially in systems such as plant-pathogen interactions where co-evolutionary signals are often shallow.

AF3 should not be interpreted as a binary predictor in plant interaction studies, but rather as a structured prioritization framework within the biological context that enriches for high-confidence candidates while still requiring integrative validation, particularly for borderline cases. The SPIN pipeline integrates bulk file inputs for protein preprocessing, structured JSON file preparation, and high-throughput AF3 inference for large-scale protein structure prediction, balancing computational efficiency with biological fidelity.

### 4.2 AlphaFold3 structural predictions for *Ca.* Liberibacter effectors and *Citrus* targets

*Ca.* Liberibacter effectors displayed highly confident *in silico* binding potential with several proteases such as S10 family serine carboxypeptidases (orange1.1g012982m and orange1.1g010909m), senescence-specific papain cysteine proteases (orange1.1g018958m and orange1.1g018104m), and M24 methionyl aminopeptidase (orange1.1g016183m). It’s conceivable that effectors may alter protein function and downstream physiological changes, thereby reprogramming the transcriptional activity observed by Franco et al. (2020) in HLB-infected citrus phloem. However, AF3 confidence scores proved inadequate to corroborate several experimentally validated interactions between Liberibacter effectors and citrus proteins (Table 4, and the references therein). Beyond structural training imbalances, multiple biological factors likely contribute to discrepancies between AF3 predictions and experimental observations.

Many PPIs occur within higher-order assemblies rather than binary complexes, often requiring additional scaffold or regulatory components (Mackey et al., 2002). Interaction specificity is further modulated by post-translational modifications such as glycosylation (Strasser, 2016) and phosphorylation (Couto and Zipfel, 2016), which are not explicitly modeled by AF3. Calcium ions (DeFalco et al., 2010), as well as membrane localization effects (Jarsch et al., 2014), introduce additional physiological context influencing protein conformation and interaction interfaces, but typically absent from the prediction workflows. Moreover, transient and weak-affinity interactions central to signaling networks are underrepresented in structural datasets, biasing models toward stable complexes (Perkins et al., 2010).

Reiterating, AF3 confidence metrics must be interpreted within both computational and biological contexts. Previous studies have indicated that an ipTM score of approximately 0.3, combined with strong intermolecular correlation signals in PAE plots, can be a practical criterion for identifying complexes suitable for detailed structural and functional assessment (Weeratunga et al., 2024). The interface-focused metrics such as ipSAE and pDockQ improve prioritization positioning AF3 as a decision-support tool rather than a substitute for empirical approaches.

### 4.3 *Ca.* Liberibacter and *Ca.* Phytoplasma citri effectors manipulate ROS detoxification enzymes and intermediary metabolism in *Citrus* phloem

SPIN-derived structural interaction datasets comparing host targets of *Ca.* Liberibacter and *Ca.* P. citri effectors revealed that, despite distinct disease phenotypes, both pathogens preferentially target antioxidant defense pathways and central carbon metabolism.

Oxidative homeostasis is a central mechanistic component of plant immunity, and successful colonization of citrus phloem depends on mitigating the pathogen-induced oxidative burden (Ma et al., 2022; Dermastia et al., 2023). AF3 prediction of *Ca.* Liberibacter effector interactions confirmed the dynamic changes in the enzymatic activities of peroxidases (PRXs), e.g., ascorbate PRX (orange1.1g025646m) and heme-dependent PRXs (orange1.1g020143m and orange1.1g018811m) discernible in citrus phloem during HLB progression (Franco et al., 2020). AF3-predicted interactions show that *Ca.* Liberibacter effectors may bind and manipulate the activities of antioxidant enzymes glutathione S-transferases (GSTs) (*τ* GST, orange1.1g027333m; □ GST, orange1.1g027910m; and Φ GST, orange1.1g046920m), glutaredoxins (orange1.1g046920m and orange1.1g027910m), and thioredoxin (orange1.1g026422m). Likewise, catalase (Ciclev10000948m), heme PRX (Ciclev10026321m), ascorbate PRXs (Ciclev10001685m and orange1.1g025646m), glutathione PRX (orange1.1g026011m), PRX5 type peroxiredoxin (orange1.1g045485m), and Φ GST (orange1.1g046920m) were predicted to be targets for *Ca.* P. citri effectors.

Liberibacters and Phytoplasmas both manipulate host cell metabolism for the provision of nutrients, energy, and metabolites for successful colonization of the plant hosts. These observations are consistent with an intracellular lifestyle of the pathogen deficient in metabolic and biosynthetic pathways for essential metabolites such as ATP, nucleotides, and essential amino acids due to extensive genome reduction (Jain et al., 2017; Duan et al., 2009; Wulff et al., 2014; Hartung et al., 2011; Cai et al., 2021; Cai et al., 2022). Similar changes were observed in primary carbon metabolic pathways and an enhanced ascorbate-glutathione cycle for relieving oxidative burden in grapevine (*Vitis vinifera*) and tobacco (*Nicotiana benthamiana*) infected with *Ca.* P. solani (ribosomal group 16SrXII-A) (Dermastia et al., 2023). Most importantly, the metabolomic changes in HLB- and Phytoplasma-infected plants (Wang et al., 2022; Tan et al., 2021) are not simply consequences of advanced disease progression but instead reflect active manipulation of host physiology. Liberibacter and Phytoplasma effectors alter key metabolic pathways, including glycolysis, the Krebs cycle, and nucleotide, fatty acid, and amino acid biosynthesis. This reprogramming of intermediary carbon metabolism facilitates colonization of the sucrose-rich yet nutritionally constrained phloem environment (Tiwari et al., manuscript in preparation).

### 4.4 *Ca.* Phytoplasma citri effectors target regulatory networks implicated in *Citrus* **growth and development**

Despite significant overlaps of protein interaction partners with Liberibacter, SPIN pipeline predicted unique protein interaction targets for *Ca.* P. citri effectors, thus supporting its ability to capture pathosystem-specific interactions. Phytoplasma infection causes aberrant morphological changes and failure of reproductive transition in citrus. High confidence predictions of molecular associations between SAP54-RAD23C/AP1/SEP3 (Fig. 4C, Table 4) is congruent with mechanistic studies in *A. thaliana* (Maclean et al., 2014). AF3 predicted likely interactions between *Ca.* P. citri effectors and *β*-actin (Ciclev10025866m), actin depolymerizing protein (orange1.1g002009m), actin-like ATPase domain protein (orange1.1g002367m), and actin-crosslinking protein (orange1.1g010600m) involved in cytoskeleton remodeling in citrus phloem. Buxa et al. (2015) had earlier observed actin-mediated tethering of *Ca.* P. solani and re-organization of sieve cell plasma membrane in infected tomato. SPIN analysis also identified citrus target proteins regulating plant architecture and development, such as the transcriptional corepressor Topless-Related Protein 1 (TRP1) (Orange1.1g001415m), and Translationally Controlled Tumor Protein (TCTP) (Ciclev10006071m). TRP1 is implicated in phytohormone and immune signaling, and maintenance of apical meristem activity (Plant et al., 2021) and TCTP is an important regulator of cell size, division and vegetative growth (Berkowitz et al., 2008). Another high confidence *Ca.* P. citri target protein was identified as CDC48 family AAA ATPase (orange1.1g003620m) has a well characterized role in plant immunity (Nicolas-Francès et al., 2024) and its inactivation causes aberrant development and early cessation of shoot and leaf growth, as well as flower sterility (Bae et al., 2009).

## Conclusion

SPIN enables systematic, structure-guided prioritization of PPIs for downstream experimental validation, thus accelerating identification of critical host vulnerabilities with potential to inform strategies for crop protection and resistance breeding. The pipeline accepts FASTA files of host and pathogen proteins as input and automates all preprocessing steps requisite for the generation of input JSON files compatible with AF3, which the user can run separately in accordance with the licensing terms of AF3. The pathogen and host preprocessing and JSON file preparation of theSPIN pipeline was wrapped as a Docker image that integrates all required dependencies (SignalP 6.0, OrthoFinder, CD-HIT) into a portable, fully reproducible environment. This containerized design enables seamless execution across diverse computing platforms without complex installation, thereby broadening accessibility for the research community.

## CRediT statement

Conceptualization, project administration, and funding acquisition: JCHT, MJ, CDM, and RL; Methodology and investigation: ZB, HK, and BD; Pipeline development: ZB; Docker imaging: HK; Data curation: ZB; Data analysis: ZB, HK, BD, MJ, MP, TT, and WB; Original manuscript draft: ZB, JCHT, MJ, WB, TT, and MP; Final review and editing: MJ, JCHT, and RL. All authors read the final version and approved the manuscript.

## Supplementary tables and data availability

Suppl. Table S0: List of 80 ground-truth positive protein interaction pairs and 324 negative (non-interacting) protein pairs.

Suppl. Table S1: Accession numbers and strain identifiers for *Ca.* Liberibacter and *Ca.* Phytoplasma citri genomes used in this study.

Suppl. Table S2: Full-length sequences of 733 *Citrus sinensis* and *C. clementina* phloem-related proteins.

Suppl. Table S3: 798 Sec-translocated proteins of *Ca.* Liberibacter.

Suppl. Table S4: 88 unique secreted proteins of *Ca.* Liberibacter.

Suppl. Table S5: 52,536 putative *Citrus*-*Ca.* Liberibacter protein interactions

Suppl. Table S6: The average computing time for predicting a protein protein interaction using SPIN.

Suppl. Table S7: AF3 native and derived confidence scores for *Citrus*-*Ca.* Liberibacter protein interactions.

Suppl. Table S8: 53 Sec-translocated proteins of *Ca.* Phytoplasma citri.

Suppl. Table S9: 14 unique sectreted proteins of *Ca.* Phytoplasma citri.

Suppl. Table S10: AF3 native and derived confidence scores for *Citrus*-*Ca.* Phytoplasma citri protein interactions.

The code for this pipeline is available at https://github.com/baozhuf/SPIN/, and Docker image is available at https://hub.docker.com/r/harshkhanna1304/spin-preprocessing

## Online resources

https://github.com/dunbracklab/IPSAE

https://alphafoldserver.com

https://services.healthtech.dtu.dk/services/SignalP-6.0/

https://www.ebi.ac.uk/interpro/search/sequence/

https://www.genome.jp/kegg/pathway.html

https://geneontology.org/

